# MYC targeted de novo binders impair tumor growth

**DOI:** 10.64898/2026.10.15.758618

**Authors:** Nora K. McNamara-Bordewick, Kejia Wu, Yulong Su, Assmaa ElSheikh, Xiaoyan Wang, Melissa Cunningham, Colin Daniel, Rosalie C. Sears, Robert N. Eisenman, David Baker

**Affiliations:** Department of Biochemistry, University of Washington, Seattle, WA, USA; Institute for Protein Design, University of Washington, Seattle, WA, USA; Division of Human Biology, Fred Hutchinson Cancer Center, Seattle, WA, USA; Division of Basic Sciences, Fred Hutchinson Cancer Center, Seattle, WA, USA; Department of Molecular and Medical Genetics, Oregon Health & Science University, Portland, OR, USA; Department of Surgery, Oregon Health & Science University, Portland, OR, USA; Brenden-Colson Center for Pancreatic Care, Oregon Health & Science University, Portland, OR, USA; Knight Cancer Institute, Oregon Health & Science University, Portland, OR, USA; Howard Hughes Medical Institute, University of Washington, Seattle, WA, USA

**Author notes:** These authors contributed equally to this work (Nora K. McNamara-Bordewick, Kejia Wu).

## Abstract

MYC is one of the most frequently dysregulated oncogenic drivers across many cancers, yet direct therapeutic targeting has remained difficult because its function is mediated by disordered regions and transient protein-protein interactions. Here we design proteins that bind epitopes within regions of MYC whose sequence is conserved between human and mouse, including intrinsically disordered motifs required for oncogenic activity. In vitro, the designs bind MYC with nanomolar affinity and in cells they engage MYC, disrupt interactions with MAX, and remodel MYC-associated protein complexes. Expression of the binders decreases cell proliferation across diverse MYC-dependent hematopoietic, liver, pancreatic, and patient-derived tumor models, and in an orthotopic pancreatic cancer model, binder-expressing tumor cells were selectively depleted. This work establishes de novo protein design as a generalizable strategy to therapeutically target intrinsically disordered transcription factors.

**One-sentence summary:** We developed a panel of de novo designed intracellular protein binders that target conserved functional regions of MYC with nanomolar affinity, remodel MYC complexes and interaction networks in cells, suppress proliferation across MYC-dependent cancer models, and are depleted from orthotopic pancreatic tumors in vivo.

## Introduction

MYC is among the most critical oncogenic drivers in human cancer^1^. Dysregulated MYC activity promotes proliferation, metabolic rewiring, immune evasion^2^, and tumor progression across a broad range of solid and hematopoietic neoplasms^3–5^. However, MYC has remained a longstanding challenge for direct therapeutic intervention. Unlike enzymes with well-defined catalytic pockets, MYC is a transcription factor that functions through intrinsically disordered regions and transient protein–protein interactions, most notably heterodimerization with MAX to regulate transcriptional programs^4,6^; its most functionally important surfaces are dynamic, extended, and only partially defined in stable conformations. Existing strategies, including indirect transcriptional suppression, pathway-level modulation, and disruption of MYC–MAX heterodimerization, have provided important proof-of-concept evidence for MYC targeting. The most advanced is the Omomyc-derived mini-protein OMO-103^7^, well tolerated in a first-in-human trial^8^, which established that interfering with MYC dimerization is therapeutically tractable. Direct targeting has nonetheless focused almost entirely on the dimerization interface, leaving the conserved regions governing MYC stability, transactivation and chromatin targeting unexplored.

Recent advances in de novo protein design now enable the direct targeting of intrinsically disordered regions^9,10^. Unlike small molecules, designed proteins engage extended surfaces rather than pockets, and we reasoned that this could provide the affinity and sequence specificity needed to target MYC directly. We hypothesized that, despite MYC’s intrinsic disorder, evolutionarily conserved sequence elements and transient structural motifs could provide sufficient geometric and sequence information for the design of compact protein binders with high affinity and specificity. We set out to design binders to conserved regions of MYC implicated in oncogenic activity, and to investigate the impact of intracellular expression of these binders on cell proliferation in multiple cancer models, including liver and pancreatic cancers.

## Results

### Functional and evolutionarily conserved epitopes of c-MYC accessible to de novo protein design

MYC is a challenging drug target because its function depends on intrinsically disordered regions and transient protein–protein interactions, features that have historically limited direct therapeutic targeting (Fig. 1A). To identify potential therapeutically-relevant MYC sites to target using de novo protein design, we analyzed sequence conservation across human and mouse MYC and mapped these conserved residues onto known functional domains. This revealed multiple conserved regions associated with MYC stabilization, transcriptional activation, repression of tumor suppressor pathways, and formation of the MYC–MAX heterodimeric transcription factor complex. Many of these regions coincide with the Myc Homology Boxes (MBs), the most highly conserved functional elements across MYC homologs and family members^11^ (Fig. 1B). We selected four target epitopes that are evolutionarily constrained, functionally important, and accessible to protein design methods. Three are intrinsically disordered loops: loop 1 in MBI, which contains the phosphodegron controlling MYC stability^12^; loop 2 in MBIIIa, which mediates transcriptional repression^13^; and loop 3 in MBIV, required for apoptosis and transformation^14^. The fourth is a helical epitope in the bHLH domain that forms the MYC–MAX dimerization and DNA-binding interface^6^ (Fig. 1B–D). Binders are named for their target epitope throughout (MYCb-L1, MYCb-L2, MYCb-L3, MYCb-H).

**Figure 1.**
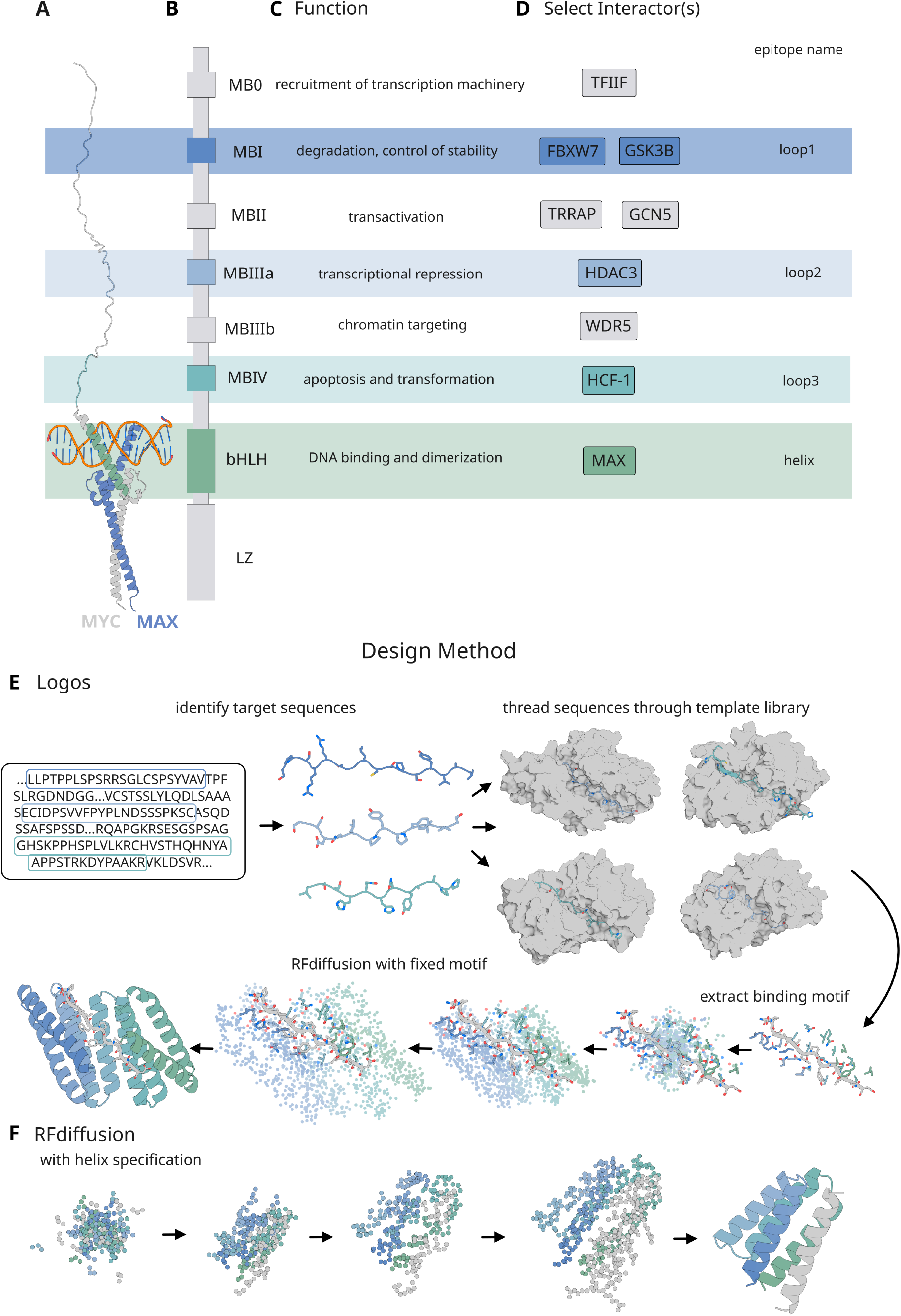
Conserved functional regions of MYC reveal targetable epitopes for de novo binder design. **(A)** Structural representation of full-length human MYC, showing the extended intrinsically disordered N-terminal and central regions and the C-terminal bHLH-LZ in complex with MAX (blue) bound to E-box DNA. Model derived from PDB 1NKP^6^ (Myc–Max bound to E-box DNA) and AlphaFold model AF-P01106 (human MYC, UniProt P01106) for the full-length disordered regions. **(B–C)** Linear domain map of MYC annotating the Myc homology boxes (MB0, MBI, MBII, MBIIIa, MBIIIb, MBIV), the bHLH, and the leucine zipper (LZ), with the putative function ascribed to each element. Shaded bands mark the four epitopes selected for design: loop 1 (MBI), loop 2 (MBIIIa), loop 3 (MBIV), and the helical epitope (bHLH). **(D)** Representative reported protein interactors for each region^11,23^. **(E)** Logos design pipeline. Target sequences spanning the selected MYC epitopes are threaded through a library of predesigned binder–peptide templates to identify compatible binding modes; the resulting binding motif is extracted and rebuilt with RFdiffusion under motif constraints, followed by sequence design and backbone refinement. **(F)** RFdiffusion pipeline with helix specification, used to generate binders against the helical bHLH epitope.

To target these four sites, we used two complementary design approaches. Logos^10^ enables recognition of intrinsically disordered sequences by threading target peptides through a diverse library of predesigned binder–peptide templates to identify compatible binding modes, followed by sequence optimization and backbone refinement (Fig. 1E). RFdiffusion^9,15^ uses a diffusion-based generative model to design de novo protein binders around structured target epitopes (Fig. 1F). Sequences were generated for designs from both methods using ProteinMPNN^16^, and those for which AlphaFold2/3^17,18^ structure predictions were close to the design models were selected for experimental characterization.

### De novo protein design yields high-affinity binders to conserved MYC epitopes

Across the four target sites, both design methods produced binders with diverse architectures surrounding their intended epitopes (Fig. 2A). The loop-targeting designs have extended backbone conformations that present linear interaction surfaces, engaging the epitope through distributed hydrogen bonding networks involving both backbone and sidechain contacts. In contrast, the helical-epitope binders have compact helical bundle architectures that cradle the target helix and achieve shape-complementary packing interactions.

**Figure 2.**
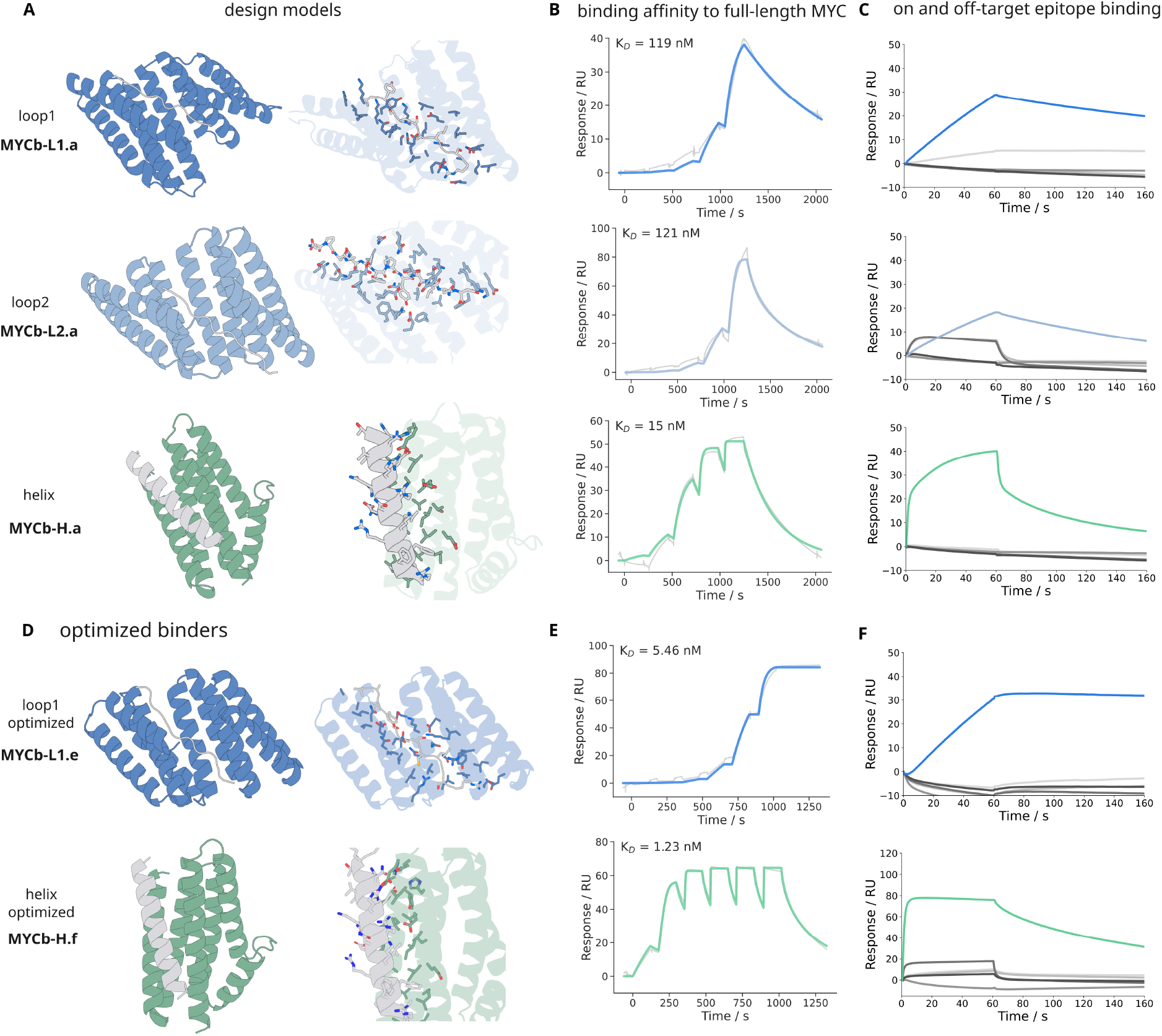
De novo design generates high-affinity, epitope-specific binders to conserved MYC regions. **(A)** Design models of representative binders to the loop 1 (MYCb-L1.a), loop 2 (MYCb-L2.a), and helical (MYCb-H.a) MYC epitopes. Left, cartoon representation of the binder (blue, loop binders; green, helix binder) with the MYC epitope in gray. Right, interface view showing sidechain contacts. **(B)** Representative SPR sensorgrams for binding of each design to full-length recombinant human MYC, with fitted K_D_ values indicated. Sensorgrams were acquired by single-cycle kinetics on a Biacore 8K: each successive peak corresponds to one injection of a 5-fold analyte series from 1.6 to 1000 nM, separated by short buffer-flow intervals, with an extended dissociation phase following the final injection. Traces are double-referenced and fitted globally to a 1:1 Langmuir binding model in Biacore Insight Evaluation Software. **(C)** SPR specificity assay. Binding of each design to its cognate MYC epitope peptide (colored) versus 3 non-cognate MYC-derived epitope peptides and 2 unrelated peptides from an orthogonal target (gray, overlaid). **(D)** Design models of affinity-optimized binders generated by motif diffusion of the loop-targeting designs and partial diffusion of the helical-epitope designs, shown as in A. **(E)** SPR sensorgrams for the optimized binders MYCb-L1.e and MYCb-H.f against full-length recombinant human MYC, acquired and fitted as in B, with fitted K_D_ values indicated. **(F)** SPR specificity of the optimized binders, shown as in C.

We expressed and purified 192 candidate designed binders and evaluated binding to full-length recombinant human MYC by surface plasmon resonance (SPR). We obtained 15 binders with affinities below 500 nM spanning three out of four targeted epitopes, including 5 binders targeting loop 1, 1 binder targeting loop 2, and 9 binders targeting the helical epitope. Representative SPR binding traces are shown in Fig. 2B for designs targeting each of the three epitopes with measured affinities ranging from 10–200 nM. To determine whether binding was driven by recognition of the intended target region, we also measured interactions with isolated on-target peptides corresponding to each design epitope (Fig. 2C). In all cases, binders retained robust binding to their cognate peptide targets, supporting the designed recognition mode.

We assessed binding specificity using a panel of non-cognate MYC-derived peptides corresponding to alternative MYC epitopes, as well as several unrelated peptides (Fig. 2C). This combined panel enabled evaluation of both within-MYC epitope selectivity and general off-target binding. The designs showed minimal detectable binding across both classes of non-cognate sequences, and any residual interactions were weak and transient with a much faster off rate compared to cognate binding, consistent with sequence- and epitope-specific recognition.

Initial designs bound full-length MYC with affinities between 10 and 200 nM, sufficient for biochemical characterization but potentially limiting for intracellular activity, where a binder must compete with endogenous partners. To improve affinity while preserving the binding modes of the validated designs, we applied two structure-guided diffusion strategies^15^ (Fig. 2D). For the loop-targeting binders, we used motif diffusion, holding the residues that contact the MYC epitope fixed in sequence and structure while rediffusing the remainder of the scaffold, thereby presenting the same interaction surface within alternative supporting architectures. For the helix-targeting binders, we used partial diffusion, applying limited noise to the parent design and denoising to sample local backbone variation around the existing binding mode without altering the overall topology. Designs were filtered using AlphaFold3^18^ and a further 174 were ordered for experimental characterization: 96 partial-diffusion designs against the helical epitope and 78 motif-diffusion designs spanning the loop 1 and loop 2 epitopes.

Optimization improved the best measured affinity for the loop epitopes from 119 nM to 5.46 nM (MYCb-L1.e) and for the helical epitope from 15 nM to 1.23 nM (MYCb-H.f) (Fig. 2E). Optimized binders retained the selectivity of their parent designs, showing minimal binding to non-cognate MYC-derived epitopes and to unrelated peptides from an orthogonal target (Fig. 2F).

### Designed binders engage MYC in cells and disrupt endogenous MYC–MAX complexes

We next evaluated whether designed binders could engage endogenous MYC in a cellular context and perturb its native interaction network. A proximity ligation assay (PLA) was used to detect proximity between MYC and either the designed binders or its endogenous obligate heterodimer MAX. Binders carried a C-terminal FLAG tag and were detected with an anti-FLAG antibody paired with an anti-MYC antibody; MYC–MAX proximity was detected by pairing anti-MYC with an anti-MAX antibody. We observed robust PLA signals between MYC and selected binders (Fig. 3A,C), indicating that de novo binders can access and engage endogenous full-length MYC in cells. Thus, the binding observed in vitro translates to target engagement in the complex intracellular environment.

**Figure 3.**
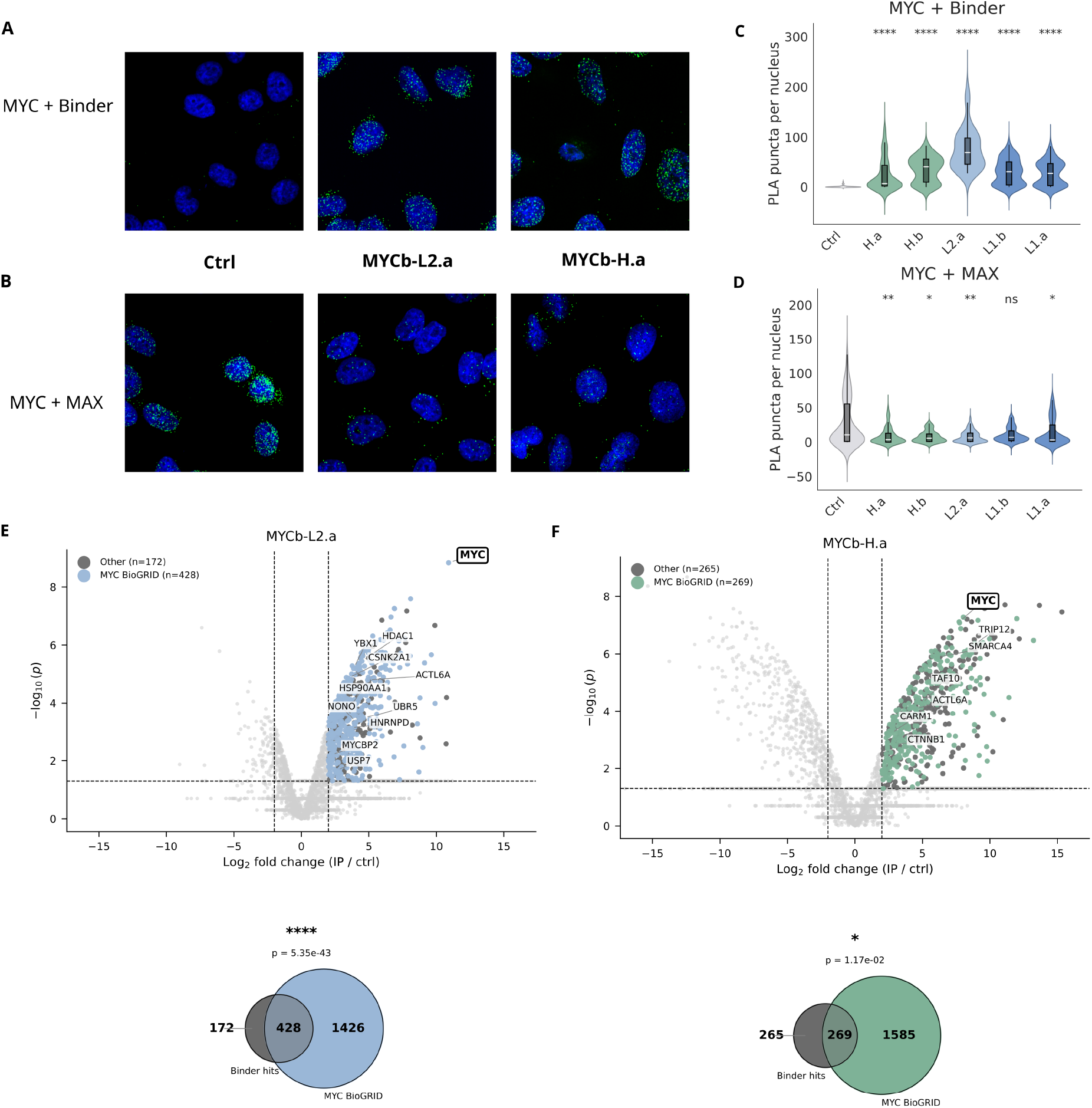
Designed binders engage MYC in cells and remodel MYC-associated complexes. **(A)** Representative proximity ligation assay (PLA) images for MYC and designed binder in HEK293T cells expressing no binder (Ctrl), MYCb-L2.a, or MYCb-H.a. PLA puncta, green; nuclei, DAPI, blue. **(B)** Representative PLA images for endogenous MYC and MAX under the same conditions, shown as in A. Binder proximity was detected with mouse anti-FLAG paired with rabbit anti-MYC; MYC–MAX proximity with rabbit anti-MYC paired with mouse anti-MAX. **(C)** Quantification of MYC–binder PLA puncta per nucleus for MYCb-H.a, MYCb-H.b, MYCb-L2.a, MYCb-L1.b, and MYCb-L1.a. Violin plots show the distribution of single-nucleus counts with median and interquartile range; fill color denotes target epitope (loop 1, loop 2, helical). Mann-Whitney U test, ****P < 0.0001 versus Ctrl. **(D)** Quantification of MYC–MAX PLA puncta per nucleus in cells expressing the same binders, shown as in C. Mann-Whitney U test; *P < 0.05, **P < 0.01; ns, not significant. **(E)** Top, volcano plot of proteins co-enriched by anti-FLAG immunoprecipitation from HEK293 cells expressing C-terminally FLAG-tagged MYCb-L2.a, relative to cells transfected with an empty FLAG-only vector. Cells were treated with 1 µM MG132 for 2 h before harvest. Proteins were quantified by DIA mass spectrometry; n = 3 replicates per condition. Significance was assessed per protein according to the replicate data available: proteins quantified in ≥2 replicates of both conditions were tested by a variance-moderated two-sample t-test on measured values only; proteins detected in one condition and entirely absent from the other were tested by Fisher’s exact test on replicate detection counts; proteins with insufficient measured replicates were not tested and are plotted as non-significant. Dashed lines, |log2 fold change| = 2 and P = 0.05. Light blue, enriched proteins annotated as MYC interactors in BioGRID^24^ (n = 428); dark gray, enriched proteins without a BioGRID MYC interaction (n = 172); light gray, proteins not passing thresholds. MYC and all curated high-confidence MYC interactors passing threshold are labeled (n = 10). Bottom, overlap between MYCb-L2.a-enriched proteins and the BioGRID MYC interactor set; Fisher’s exact test (one-sided), P = 5.35 × 10^−43^. Background for the enrichment test comprises the 4,055 proteins quantified in both IP-MS experiments (1,854 of them BioGRID MYC interactors); this background is shared between E and F so the two panels are directly comparable. **(F)** As in E, for MYCb-H.a, with enriched BioGRID interactors in green (n = 269) and other enriched proteins in dark gray (n = 265); 6 curated high-confidence interactors labeled. Overlap with the BioGRID MYC interactor set, P = 1.17 × 10^−2^.

We next assessed the effects of the designs on MYC interaction with its partner MAX (Fig. 3B,D). Because MYC–MAX heterodimerization is required for transcriptional activity, MYC– MAX proximity serves as a readout of oncogenic complex formation. Expression of four of the five binders resulted in a marked reduction in MYC–MAX PLA signal relative to controls. For MYCb-H.a this was expected, because its epitope lies within the bHLH dimerization interface. For the loop-targeting binders, however, this was surprising because N-terminal MYC interactions are not known to regulate MYC-MAX dimerization^6,11^. One possibility is that the binders compete with native interaction partners: many characterized MYC interactors contact both N- and C-terminal regions^4,11^, so engagement of a disordered N-terminal loop could disrupt complexes spanning both domains and perturb the chromatin-associated pool of MYC in which MYC–MAX complexes reside. Reduced antibody accessibility within the binder-bound complex could also contribute, since PLA reports both proximity and epitope availability.

To further characterize the consequences of MYC engagement, we expressed FLAG-tagged binders in HEK293 cells, immunoprecipitated each binder with anti-FLAG resin, and identified co-enriched proteins by data-independent acquisition mass spectrometry; enrichment is reported relative to cells transfected with an empty FLAG-only vector. We performed IP-MS for one loop-targeting and one helix-targeting design. The loop 2 binder MYCb-L2.a showed robust enrichment of endogenous MYC, with MYC ranking among the most highly enriched proteins in the pulldown (Fig. 3E). This binder also co-enriched many established MYC-associated proteins, including the histone deacetylase HDAC1, the chromatin-remodeling subunit ACTL6A, the ubiquitin ligases MYCBP2 and UBR5, and the deubiquitinase USP7, consistent with capture of endogenous MYC-containing assemblies.

The helix-targeting binder, MYCb-H.a, also strongly enriched endogenous MYC in the IP-MS (Fig. 3F). This design co-enriched a partially overlapping but distinct subset of published MYC-associated proteins, including the chromatin remodeler SMARCA4, the coactivators CARM1 and TAF10, and CTNNB1, suggesting engagement of MYC within a different subset of regulatory or chromatin-associated complexes. Across both binders, the presence of canonical MYC interactors supports endogenous engagement of MYC in cells, while differences in co-enriched partners are consistent with epitope-dependent remodeling of MYC-associated assemblies. Additional proteins detected in each dataset likely reflect context-dependent or indirect associations and will require further mechanistic validation.

Together, these results demonstrate that de novo designed binders engage endogenous MYC in cells, interfere with MYC-MAX interactions and differentially pull down relevant MYC interacting proteins depending on the targeted epitope.

### MYC binders suppress proliferation across diverse MYC-dependent cancer models

After establishing that the designed binders engage endogenous MYC and disrupt MYC–MAX complexes, we next asked whether these molecular effects translate into suppression of cancer cell growth. To enable parallel evaluation of multiple designs, we generated a lentiviral library of uniquely barcoded MYC binders that performed best in our in vitro SPR assays. Using this lentiviral library, we performed pooled competitive growth assays in MYC-dependent cancer cell lines. Following transduction, changes in binder abundance were quantified by barcode sequencing after seven days of growth, with depletion serving as a measure of impaired cellular proliferation (Fig. 4A).

**Figure 4.**
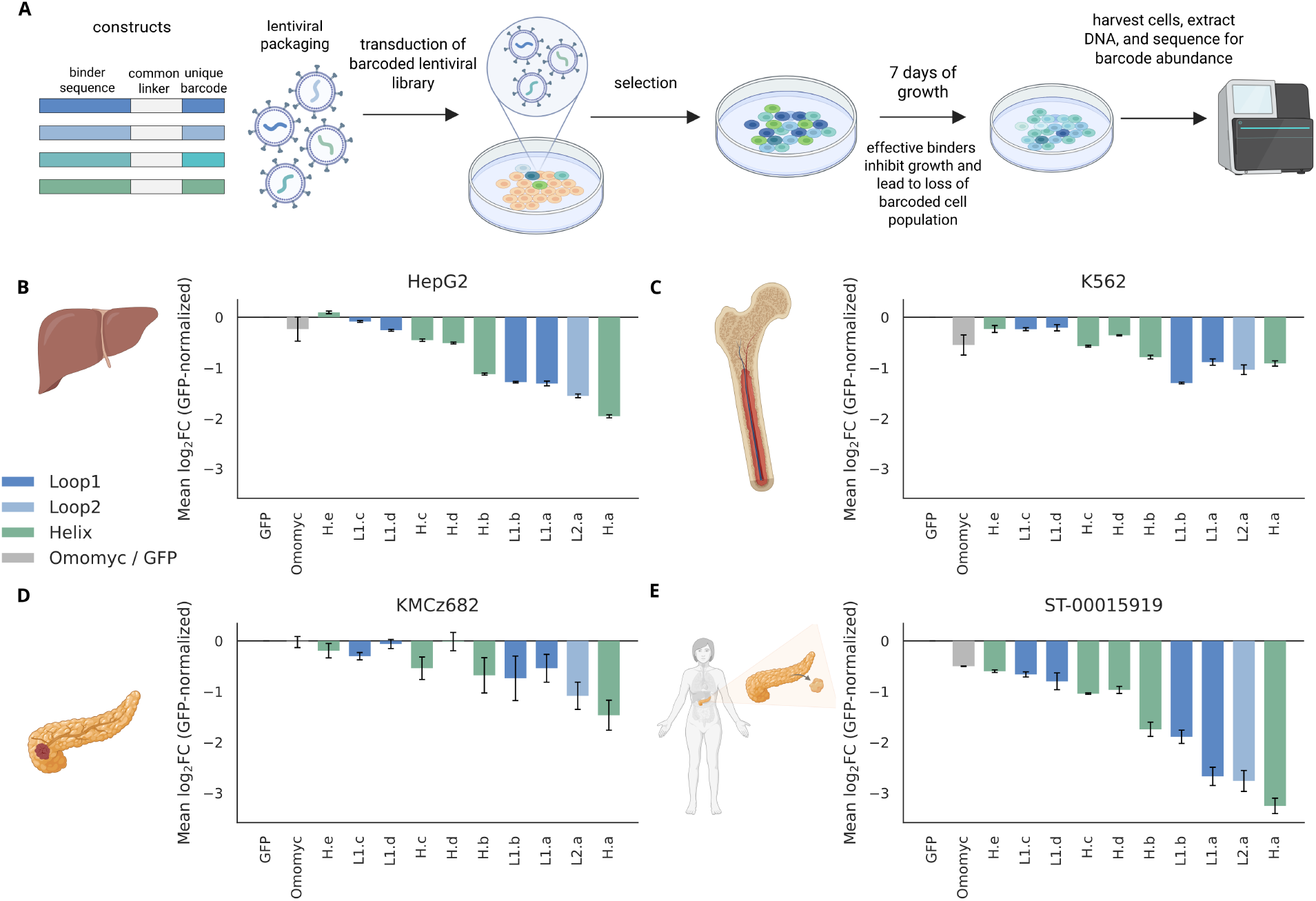
De novo MYC binders inhibit proliferation across multiple MYC-dependent cancer models. **(A)** Schematic of the pooled barcoded competitive growth assay. Ten MYC binders (MYCb-L1.a–d, MYCb-L2.a, MYCb-H.a–e), Omomyc, and GFP were cloned as barcoded constructs, packaged into lentivirus, and transduced as a pool; after selection and 7 days of growth, genomic DNA was extracted and barcode abundance quantified by amplicon sequencing. Depletion of a barcode relative to the input pool indicates impaired proliferation of cells expressing that construct. **(B)** Mean log2 fold change in barcode abundance after 7 days in HepG2 hepatoblastoma cells, normalized to GFP. Bars, mean of n = 3 independent infections; error bars, SEM. Bar color denotes target epitope: dark blue, loop 1; light blue, loop 2; green, helical. Gray, Omomyc and GFP controls. Constructs are shown in a fixed order across B–E, in ascending order of mean log2 fold change in HepG2, with controls at left. **(C)** As in B, in K562 chronic myelogenous leukemia cells. **(D)** As in B, in KMCz682 murine pancreatic ductal adenocarcinoma cells. **(E)** As in B, in the patient-derived pancreatic cancer line ST-00015919.

Across MYC-dependent cancer models spanning human liver cancer (HepG2^19^, Fig. 4B), human chronic myelogenous leukemia (K562, Fig. 4C), and murine pancreatic ductal adenocarcinoma (KMCz682^20^, Fig. 4D), nine of ten designs depleted relative to GFP in all four lines. The same five designs (MYCb-H.a, MYCb-L2.a, MYCb-L1.a, MYCb-L1.b and MYCb-H.b) were the five most strongly depleted in every model tested. These five span all three targeted epitopes, indicating that engagement at the disordered MBI and MBIIIa loops impairs proliferation as effectively as engagement at the bHLH dimerization interface. Depletion of these five designs exceeded that of Omomyc in every line under matched assay conditions, and the strongest design exceeded it by 2.4- to 62-fold (Fig. 4B–E). Thus de novo designed binders can perturb MYC-dependent growth programs.

To assess activity in more clinically relevant settings, we evaluated active binders in a patient-derived cell line generated from a primary resected tumor of a PDAC patient (Fig. 4E). Similar growth inhibitory effects were observed in this model, indicating that the consequences of MYC engagement are conserved across genetically and biologically distinct MYC-dependent cancers.

### De novo MYC binders suppress tumor growth in vivo in orthotopic pancreatic cancer models

Finally, we evaluated the in vivo activity and therapeutic potential of de novo MYC binders in an orthotopic pancreatic cancer model. To determine whether MYC engagement is sufficient to impair tumor growth in vivo, we transduced murine PDAC KMCz682 cells, derived from autochthonous tumors of the KMC genetically engineered mouse model (*Ptf1a-Cre*^*ERT2*^; *LSL-Kras*^*G12D/+*^; *Rosa-LSL*^*Myc/Myc*^) in immunocompetent C57BL/6 mice^20^, with a pooled lentiviral library of barcoded MYC binders. On day zero, a portion of the transduced cells was harvested for genomic DNA extraction to establish baseline barcode abundance, and 50,000 transduced cells were implanted orthotopically into the pancreas of syngeneic recipient mice. After fourteen days of tumor growth, tumors were harvested and binder abundance was quantified by barcode sequencing (Fig. 5A). Consistent with the competitive growth assays performed in cultured cells, active MYC binders were selectively depleted in vivo (Fig. 5B). These results demonstrate that intracellular binding of conserved and functionally important MYC epitopes is sufficient to suppress tumor growth in an in vivo cancer model.

**Figure 5.**
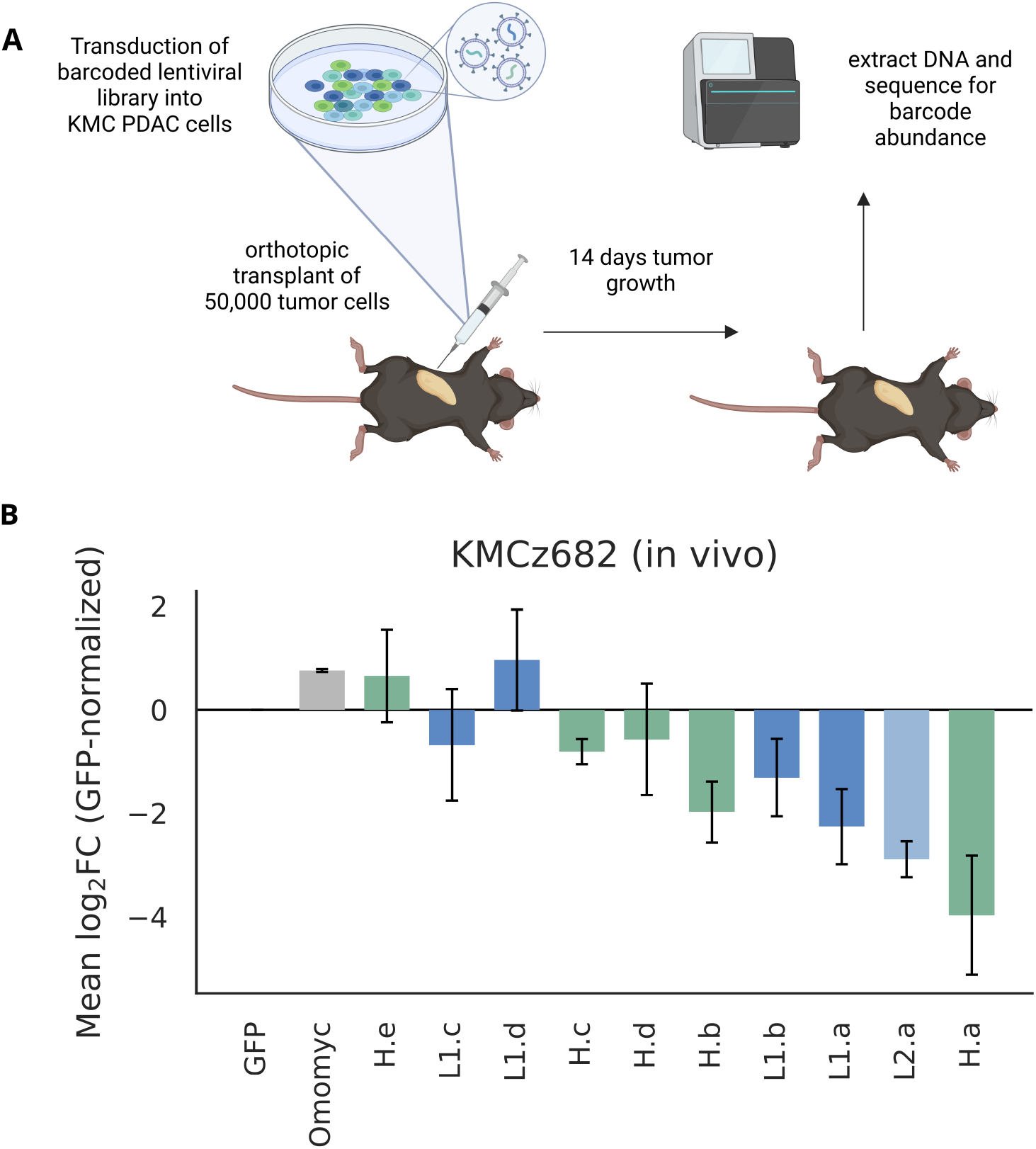
De novo MYC binders are depleted from orthotopic pancreatic tumors in vivo. **(A)** Schematic of the in vivo pooled screen. KMCz682 PDAC cells transduced with the barcoded binder library were transplanted orthotopically (50,000 cells) into the pancreas of 10-week-old male C57BL/6 mice; after 14 days of tumor growth, tumors were harvested and barcode abundance quantified by amplicon sequencing. **(B)** Mean log2 fold change in barcode abundance in orthotopic KMCz682 tumors relative to the day-zero input cell pool, normalized to GFP. Bars, mean of n = 2 sequencing samples, each pooled from 2 tumors (4 mice total), relative to n = 3 day-zero input samples; error bars, SEM. Bar color denotes target epitope: dark blue, loop 1; light blue, loop 2; green, helical. Gray, Omomyc and GFP controls. Constructs are shown in the same order as Fig. 4B–E.

## Discussion

This study demonstrates that de novo protein design can target MYC, a historically “undruggable” cancer driver. By targeting multiple conserved regions of MYC, we identified binders that not only show strong biochemical binding, but also produce consistent functional effects in cells. Across MYC-dependent cancer models, including blood, liver, pancreatic, and patient-derived systems, the five most active binders reduced cancer cell growth and outperformed Omomyc^7^ under matched conditions. In vivo experiments further demonstrate that the designed binders suppress MYC-driven proliferation, as active binders deplete tumor cells in an orthotopic pancreatic cancer mouse model, consistent with reduced tumor fitness after MYC engagement. Taken together, these results suggest a path from designed intracellular binders to therapeutically-relevant MYC inhibition in animal models and eventually in cancers where MYC remains a major “undruggable” target.

More broadly, this work points toward a new route for therapeutically targeting intracellular proteins that have been difficult to address with small molecules or antibodies. Many of the most important disease drivers, including transcription factors, viral proteins, fusion proteins, and neurodegeneration-associated proteins, contain intrinsically disordered regions, transient interaction motifs, or short conserved sequence elements rather than stable pockets. De novo protein design offers a way to recognize these features directly and build new therapeutic molecules around them.

Critical to therapeutic application of our MYC inhibiting designs, and other de novo designs targeting intracellular proteins, is delivery into the cell. A promising route to achieving this is LNP-mRNA delivery^21^, particularly for targets such as MYC in tissues such as the liver which readily take up LNPs through ApoE adsorption and LDL receptor-mediated hepatocyte uptake^22^. The combination of programmable protein design and nucleic-acid delivery could open a broader class of intracellular biologic therapeutics for MYC-driven cancer, viral disease, and other disorders caused by disordered or previously inaccessible proteins.

## Methods

### MYC sequence analysis and selection of target epitopes

Human MYC (UniProt P01106) and mouse MYC protein sequences were compared to identify conserved regions associated with established MYC regulatory functions. Published annotations of the MYC homology boxes, transcriptional regulatory regions and C-terminal basic helix– loop–helix leucine-zipper (bHLH-LZ) domain were used to guide target selection^6,11–14^. Four segments were selected for binder design: loop 1 within MYC homology box I (MBI), which contains a phosphodegron linked to MYC turnover; loop 2 within MBIIIa; loop 3 within MBIV; and a helical region of the bHLH domain that participates in MYC–MAX association. The targeted segments were human MYC residues 55–77 (loop 1, MBI), 187–208 (loop 2, MBIIIa), 286–324 (loop 3, MBIV) and 356–379 (helical epitope, bHLH), numbered according to UniProt P01106.

The structural illustration of MYC in Fig. 1 combines the MYC–MAX heterodimer bound to E-box DNA (PDB 1NKP) with the predicted full-length human MYC model (AlphaFold model AF-P01106). This composite was used to illustrate the relative positions of annotated MYC domains and the selected epitopes, rather than to represent a uniquely determined structure of full-length MYC.

### Design of binders to disordered MYC epitopes

Binders recognizing the three loop epitopes were generated using the Logos sequence-directed peptide-binder design framework^10^. In this approach, each MYC epitope sequence was threaded through an existing collection of computationally constructed binder–peptide complexes representing diverse extended peptide geometries. Candidate starting models were assessed for compatibility between the target residues and the amino-acid recognition pockets of the template. Interface fragments providing favorable contacts to the target peptide were retained as binding motifs, and RFdiffusion was used to construct supporting protein backbones around these motifs^10,15^. ProteinMPNN was then applied to generate binder sequences compatible with the resulting structures^16^. Designs underwent iterative sequence and structural refinement before evaluation with protein-complex structure prediction. Predicted complexes were filtered using AlphaFold-derived confidence metrics (pLDDT of binder > 90 and interface PAE < 10); passing designs were clustered by sequence diversity and representatives were selected by visual inspection for experimental testing. The template library was the same as that described previously^10^. The threading and scoring settings, diffusion parameters and sequence-design settings were the same as those previously reported^10^.

### Design of binders to the MYC helical epitope

The helical bHLH epitope was targeted using RFdiffusion-based binder generation with the intended target region represented in a helical conformation^9,15^. The model was used to sample binder backbones surrounding the specified epitope, after which ProteinMPNN generated sequences for the proposed binder scaffolds^16^. Predicted binder–target complexes were examined using AlphaFold2 and/or AlphaFold3^17,18^ to assess whether the proposed interaction geometry and binder fold were recovered. Across the four target regions, 192 initial candidates were selected for experimental testing as above.

### Structure-guided affinity optimization

Experimentally validated parent designs were subjected to a second round of backbone diversification intended to improve binding while retaining their original epitope-recognition modes. For binders to loop 1 and loop 2, contact residues at the designed peptide interface were retained in both sequence and geometry while RFdiffusion rebuilt or remodeled the remaining supporting scaffold (motif-constrained diffusion). For binders recognizing the helical epitope, partial diffusion was used to introduce limited structural variation into the starting binder model, followed by denoising to sample related backbone conformations. New sequences were designed with ProteinMPNN and the complexes were evaluated with AlphaFold2 and/or AlphaFold3 as above. Ninety-six helix-targeting designs (partial diffusion) and 78 loop-targeting designs (motif-constrained diffusion) were chosen for the second experimental round (174 additional candidates in total).

### Recombinant MYC binders and peptide reagents

DNA sequences encoding the computationally designed MYC binders were optimized for bacterial expression and synthesized as double-stranded DNA fragments. The genes were assembled by Golden Gate cloning into the LM0627 bacterial expression vector, which encodes a C-terminal hexahistidine tag and transformed into *Escherichia coli* BL21(DE3), following previously established protein-production procedures^9,10^.

For initial screening, individual transformants were grown overnight in LB medium containing kanamycin and transferred into Terrific Broth-based autoinduction medium supplemented with kanamycin, 2 mM MgSO_4_ and 1× 5052 autoinduction mixture. Cultures were incubated for approximately 24 h, initially at 37 °C for 6 h followed by 30 °C for 18 h. Cells were harvested by centrifugation and lysed using BugBuster reagent supplemented with DNase I (0.01 mg ml^−1^) and protease inhibitors. Clarified lysates were subjected to immobilized metal affinity chromatography using Ni-NTA agarose resin. The resin was washed with 25 mM Tris-HCl (pH 8.0), 300 mM NaCl and 30 mM imidazole, and the bound proteins were eluted using the same buffer containing 400 mM imidazole.

For further characterization, selected binders were expressed in 50-ml autoinduction cultures, grown at 37 °C for the first 6 h and at 22 °C for approximately 18 h. Cell pellets were resuspended in lysis buffer containing 25 mM Tris-HCl, 150 mM NaCl, 0.1 mg ml^−1^ lysozyme, mg ml^−1^ DNase I, 1 mM PMSF and protease inhibitors. Following sonication and clarification by centrifugation, soluble proteins were purified using Ni-NTA agarose resin and eluted with an imidazole-containing buffer. Selected proteins were further purified by size-exclusion chromatography on a Superdex 75 Increase 10/300 GL column (Cytiva) equilibrated with 25 mM Tris-HCl (pH 8.0) and 150 mM NaCl. Fractions containing the desired protein were collected, and concentrations were determined by absorbance at 280 nm using sequence-derived extinction coefficients.

Full-length recombinant human MYC (residues 1–439, UniProt P01106) was purchased from CUSABIO (catalog number CSB-EP015270HU-B), expressed in *E. coli* with an N-terminal MBP tag and a C-terminal 6xHis-Avi tag and supplied biotinylated, and was used for SPR characterization of the designed binders. The four targeted MYC epitopes (loop 1, loop 2, loop 3 and the helical epitope) and two unrelated epitopes were expressed in *E. coli* as fusions to a SUMO solubility scaffold carrying an N-terminal AviTag, followed by a PAAGS linker and the epitope, and the AviTag was biotinylated in vitro by BirA.

### Surface plasmon resonance and binding analysis

Binding experiments were performed by surface plasmon resonance (SPR) on a Biacore 8K instrument (Cytiva) in HBS-EP+ running buffer supplemented with 1 mM TCEP. Biotinylated full-length MYC or the biotinylated AviTag-SUMO epitope fusions were captured on a Series S Sensor Chip CAP using the Biotin CAPture Kit (Cytiva). For representative affinity measurements, the initial designs were injected in a five-point, fivefold dilution series spanning 1.6–1,000 nM (1.6, 8, 40, 200 and 1,000 nM) using a single-cycle kinetic protocol, and the optimized designs were injected in a six-point, fivefold dilution series spanning 1.6–5,000 nM (1.6, 8, 40, 200, 1,000 and 5,000 nM). Successive injections were separated by brief buffer-flow periods, followed by an extended dissociation phase after the final injection. Sensorgrams were double-referenced and globally fitted to a 1:1 Langmuir interaction model using Biacore Insight Evaluation Software. Equilibrium dissociation constants (*K*_D_) were obtained from the fitted association and dissociation rate constants.

For specificity measurements, each design was examined at 1000 nM against its cognate MYC-derived peptide, three peptides representing alternative MYC epitopes and two unrelated peptides from an orthogonal target. The same general strategy was applied to optimized designs.

### Cell lines and culture conditions

HEK293T cells were used for proximity ligation assays, and HEK293 cells were used for binder affinity purification followed by mass spectrometry. The pooled competitive growth assays used HepG2 cells (a human hepatoblastoma-derived line), K562 human chronic myelogenous leukemia cells, KMCz682 murine pancreatic ductal adenocarcinoma (PDAC) cells and the patient-derived PDAC line ST-00015919. KMCz682 cells were derived from the KMC genetically engineered mouse model (*Ptf1a-Cre*^ERT2^; *LSL-Kras*^G12D/+^; *Rosa-LSL*^Myc/Myc^), as described previously^20^. ST-00015919 was derived from a primary resected human PDAC tumor.

### In situ proximity ligation assays

MYC binders carrying a C-terminal FLAG tag were expressed in HEK293T cells, and proximity to endogenous MYC was assessed by in situ proximity ligation assay (PLA). MYC–binder proximity was detected using mouse anti-FLAG and rabbit anti-MYC primary antibodies. In parallel, endogenous MYC–MAX proximity was measured using rabbit anti-MYC and mouse anti-MAX antibodies. Cells without binder expression served as controls. After PLA signal development, nuclei were counterstained with DAPI.

PLA puncta were quantified on a per-nucleus basis from microscopy images. Single-nucleus puncta counts were displayed as distributions, with medians and interquartile ranges. Comparisons between each binder and the no-binder control were performed using the two-sided Mann–Whitney U test, as indicated in Fig. 3.

### Anti-FLAG immunoprecipitation and mass spectrometry

Binder coding sequences were cloned into pcDNA3 with a C-terminal FLAG tag, and an empty vector encoding the FLAG tag alone served as the reference condition. HEK293 cells were grown to sub-confluence and transfected with 1 µg of plasmid per 1 million cells to express MYCb-L2.a or MYCb-H.a, or with an empty FLAG-only vector as the reference condition. Before harvest, cells were treated with 1 µM MG132 for 2 h. Protein complexes associated with the expressed binders were isolated using anti-FLAG affinity resin. Three biological replicates were analyzed for each condition.

### Proteomics statistical analysis and MYC-interactor annotation

Enrichment was assessed separately for MYCb-L2.a and MYCb-H.a relative to the FLAG-only control. For proteins quantified in at least two replicates in both groups, measured protein quantities were compared by a variance-moderated two-sample t-test without substituting values for missing measurements. Proteins detected in one group but absent from all replicates of the other group were evaluated using Fisher’s exact test on replicate-level detection counts. Proteins with insufficient observations for either analysis were not assigned a significance test. Proteins were displayed according to their log_2_ enrichment and statistical significance, with visualization thresholds of |log_2_ fold change| = 2 and P = 0.05.

Published MYC-associated proteins were annotated using BioGRID^24^. For analyses of interactor enrichment, the background was restricted to the 4,055 proteins quantified in both IP–MS experiments, of which 1,854 were annotated MYC interactors in the dataset used for the analysis. The intersection between proteins enriched by each binder and BioGRID-annotated MYC interactors was evaluated using a one-sided Fisher’s exact test.

### Pooled lentiviral competitive growth screens

Ten MYC-binding designs (MYCb-L1.a–d, MYCb-L2.a and MYCb-H.a–e), together with Omomyc and GFP controls, were individually assigned identifying DNA barcodes and assembled into a pooled lentiviral expression library. The library was packaged and used to transduce HepG2, K562, KMCz682 and ST-00015919 cells. Following selection (d0), transduced cultures were maintained for seven to fourteen days, after which genomic DNA was isolated and the integrated barcodes were quantified by amplicon sequencing. Three independent infections were analyzed for each cell model.

For each construct, changes in barcode abundance between the input reference and the end of the growth interval were expressed as log_2_ fold changes and normalized to the GFP control within the corresponding experiment. Mean normalized log_2_ fold changes and standard errors of the mean were calculated across the three independent infections.

### Orthotopic pancreatic tumor competitive fitness assay

KMC z682 murine PDAC cells carrying the pooled barcoded lentiviral binder library were used to evaluate the representation of individual constructs during tumor growth in vivo. Cells were transduced at a low multiplicity of infection (MOI) to ensure that the majority of cells incorporated a single barcode, followed by selection with puromycin. After selection, cells were allowed to recover and expand for two additional days; the resulting population was designated day 0 (D0). An aliquot of this transduced cell pool was collected at D0 for genomic DNA extraction to establish baseline barcode abundance (three input sequencing samples). For each recipient, 50,000 transduced cells were implanted orthotopically into the pancreas of a 10-week-old male syngeneic C57BL/6 mouse. Tumors were harvested 14 days after implantation. Four mice were included; the harvested tumors were combined into two sequencing samples, each comprising material from two tumors. Genomic DNA was extracted, and barcode amplicons were sequenced by next-generation sequencing.

### Statistical analysis and reproducibility

The experimental unit and statistical analysis were selected according to assay type. SPR sensorgrams were analyzed by global fitting to a 1:1 kinetic binding model. Single-cell PLA puncta distributions were compared with the no-binder control using the Mann–Whitney U test. IP–MS differential enrichment was analyzed using variance-moderated tests of measured protein abundance or Fisher’s exact tests of detection counts, according to data availability, and enrichment of known MYC interactors was evaluated by a one-sided Fisher’s exact test. In pooled cell growth assays, summary values represent the mean ± SEM from three independent infections. For the orthotopic experiment, the two sequenced tumor pools each contained material from two mice; these pools, rather than individual mice, are the units represented by the error bars. Where no statistical significance test was performed, observed differences are described without assigning a P value.

## Acknowledgments

This work was funded by the National Institutes of Health (National Institute of Allergy and Infectious Diseases grant R01AI160052; National Cancer Institute grants R01CA312340 and R35CA231989 to R.N.E., and R01CA186241 and U01CA294548 to R.C.S.), the Gates Foundation (grant INV-043758), and the Howard Hughes Medical Institute (D.B.).

## Author Contributions

N.K.M.-B., K.W., and D.B. conceptualized the study. K.W. developed the Logos platform used for disordered-epitope design. N.K.M.-B. and K.W. designed the MYC binders, performed the structure-guided optimization, and selected designs for experimental testing. N.K.M.-B. expressed and purified the binders and performed SPR. Y.S. performed the proximity ligation assays and the immunoprecipitation and mass spectrometry experiments. Y.S. generated the barcoded lentiviral library and performed the pooled proliferation screens. A.E. performed the pooled screen in the patient-derived line and performed the orthotopic in vivo screens, with assistance from X.W., M.C., and C.D. R.C.S. and R.N.E. supervised the cell biology and in vivo experiments. D.B. supervised the research. N.K.M.-B. wrote the manuscript with input from all authors. All authors reviewed and commented on the manuscript.

## Data Availability

The data supporting the findings of this study are provided in the main text and associated data repositories. Amino acid sequences and computational design models of the MYC binders, together with design and analysis scripts and numerical source data underlying the figures, will be deposited prior to journal publication. Raw and processed mass spectrometry proteomics data will be deposited prior to journal publication. Barcode sequencing data from pooled cell proliferation and in vivo tumor experiments, together with processed barcode counts and associated analysis scripts, will be made available prior to journal publication. Plasmids encoding the designed MYC binders will be available from the corresponding author upon reasonable request, subject to applicable institutional material transfer agreements.

